# Economic performance and cost-effectiveness of using a DEC-salt social enterprise for eliminating the major neglected tropical disease, lymphatic filariasis

**DOI:** 10.1101/502542

**Authors:** Swarnali Sharma, Morgan E. Smith, James Reimer, David B. O’Brien, Jean M. Brissau, Marie C. Donahue, Clarence E. Carter, Edwin Michael

**Affiliations:** Department of Biological Sciences, University of Notre Dame, Galvin Life Science Center, Notre Dame, IN 46556, USA; 1071 Devonshire Road, Grosse Pointe Park, MI 48230, USA; 10720 Serenbe Lane, Chattahoochee Hills, GA 30268, USA; College of Science, University of Notre Dame, Notre Dame, IN 46556, USA; Eck Institute of Global Health, University of Notre Dame, 305 Brownson Hall, Notre Dame, IN 46556, USA; College of Science, 215 Jordon Hall of Science, University of Notre Dame, Notre Dame, IN 46556, USA

## Abstract

**Background:** Salt fortified with the drug, diethylcarbamazine (DEC), and introduced into a competitive market has the potential to overcome the obstacles associated with tablet-based Lymphatic Filariasis (LF) elimination programs. Questions remain, however, regarding the economic viability, production capacity, and effectiveness of this strategy as a sustainable means to bring about LF elimination in resource poor settings.

**Methodology and Principal Findings:** We evaluated the performance and effectiveness of a novel social enterprise-based approach developed and tested in Léogâne, Haiti, as a strategy to sustainably and cost-efficiently distribute DEC-medicated salt into a competitive market at quantities sufficient to bring about the elimination of LF. We undertook a cost-revenue analysis to evaluate the production capability and financial feasibility of the developed DEC salt social enterprise, and a modeling study centered on applying a dynamic mathematical model localized to reflect local LF transmission dynamics to evaluate the cost-effectiveness of using this intervention versus standard annual Mass Drug Administration (MDA) for eliminating LF in Léogâne. We show that the salt enterprise because of its mixed product business strategy may have already reached the production capacity for delivering sufficient quantities of edible DEC-medicated salt to bring about LF transmission in the Léogâne study setting. Due to increasing revenues obtained from the sale of DEC salt over time, expansion of its delivery in the population, and greater cumulative impact on the survival of worms, this strategy could also represent a significantly more cost-effective option than annual DEC tablet-based MDA for accomplishing LF elimination.

**Significance:** A social enterprise approach can offer an innovative market-based strategy by which edible salt fortified with DEC could be distributed to communities both on a financially sustainable basis and at sufficient quantity to eliminate LF. Deployment of similarly fashioned intervention strategies would improve current efforts to successfully accomplish the goal of LF elimination, particularly in difficult-to-control settings.

**Author summary:** With less than three years remaining for meeting the 2020 target set by WHO for accomplishing the global elimination of Lymphatic Filariasis (LF), concerns are emerging regarding the feasibility of meeting this goal using the current tablet-based Mass Drug Administration strategy. Salt fortified with the antifilarial drug, diethylcarbamazine (DEC), could offer an intervention that avoids many of the barriers connected with tablet-based elimination programs. We analyzed the economic performance and cost-effectiveness of a novel DEC-salt social enterprise developed and tested in Léogâne arrondissement, Haiti, as a particularly significant strategy for accomplishing sustainable LF elimination in such complex settings. We show that because of increasing revenue from the sale of the DEC salt over time, expansion of its delivery in the population, and the adverse effect of continuous consumption of the drug on worms, the delivery of DEC through a salt enterprise can represent a significantly more cost-effective option than annual DEC tablet-based MDA for accomplishing LF elimination in settings, like Léogâne. We indicate that development of policy and research into how to deploy similarly-fashioned interventions, or work with the salt industry to increase population use of medicated salt, would improve present efforts to successfully accomplish the elimination of LF.

## Introduction

Lymphatic filariasis (LF), a mosquito-borne neglected tropical disease (NTD), commonly known as elephantiasis, is one of only parasitic six diseases currently targeted for potential global eradication by 2020 using preventive mass chemotherapy [1–3]. Despite the impressive expansion of a WHO-led elimination program aimed toward the meeting of this goal in all endemic countries since 2000, stakeholders committed to global LF elimination have recognized that the current tablet-based mass drug intervention is resource-intensive, can face significant compliance issues with time, and may be difficult to implement in remote or socio-ecologically complex areas, such as urban and socio-politically unstable settings, hampering foreseen elimination goals [4–8]. These difficulties have heightened interest in investigating the impacts of either approaches aimed at scaling-up treatment strategies or inclusion of preventive activities into drug programs (such as supplemental vector control), or evaluation of novel intervention technologies, that can effectively overcome current barriers in order to accelerate parasite elimination [9–11].

Salt fortified with the anti-filarial drug, diethylcarbamazine (DEC), could offer an intervention that avoids many of the above issues connected with tablet-based elimination programs [18]. Indeed, DEC-salt has played a major role in the elimination of LF in a number of pilot and region-wide settings in Africa, Central America, and Asia [5,12–17]. The low dose of DEC (0.1-0.6% [w/w]) used in these studies and programs was well tolerated and rarely associated with adverse reactions. It also has the potential to be more effective than tablet-based Mass Drug Administration (MDA) programs via reduction of the durations of intervention required to interrupt parasite transmission [18]. Moreover, fortified salt can also be provided to a population without developing a dedicated public distribution system, overcoming the need for developing an effective health infrastructure capable of distributing anti-filarial drugs at the high coverages needed for achieving elimination [5,7,8].

Haiti is one of only four countries remaining in the Americas where LF is still endemic [8,19]. MDA using DEC first started in the country under the National Program to Eliminate LF (NPELF) in 2000, and following funding, sociopolitical and natural disaster-based challenges to scaling up, the program realized full national coverage by 2012 [5,7,8,20]. This delay together with the technical challenge of interrupting transmission in areas of highest prevalence even with good levels of coverage using the suggested five successive years of annual MDA [5,7,8,20], indicate that it is unlikely that NPELF will meet the goal of accomplishing LF elimination in the country by the target year of 2020. In 2006, partly to overcome the above issues, a project was initiated with the collaboration of Congregation de Sainte Croix, the Notre Dame Haiti Program, and the Ministry of Public Health and the Population (MSPP), focused on the local processing and marketing of DEC-mediated salt co-fortified with potassium iodate as an alternative means to facilitate the elimination of LF (and prevention of iodine deficiency disorders) in Haiti [8]. Based on the principle of employing a social enterprise framework for providing goods and services in an entrepreneurial and innovative fashion to solve social problems [21–23], this project has since upgraded its salt production facility to facilitate a transition to the production and sale of processed food-grade co-fortified salt, raising the potential of using DEC-medicated salt as a long-term sustainable means to accomplish LF elimination in settings, such as Haiti.

Here, our major aim was to examine the economic performance and effectiveness of using the Haitian social enterprise-based framework for producing and marketing DEC-fortified salt as a sustainable, cost-effective, model for achieving the long-term elimination of LF, focusing on Léogâne arrondissement, Haiti. A cost-revenue analysis combined with a mathematical modeling-based evaluation of the cost-effectiveness of the DEC salt social enterprise compared to standard MDA was carried out to undertake this analysis. Specifically, we evaluated the economic performance and social value of the enterprise by assessing: 1) the growth in salt production, costs of resources consumed, and revenues from sales gained to determine break-even points, 2) the impact of the product-mix used for realizing the socially-relevant sale price of the salt, and 3) its cost-effectiveness compared to tablet-based MDA for accomplishing LF elimination in the study setting of Léogâne arrondissement, Haiti. We discuss the results in terms of how using a social enterprise can offer a sustainable and innovative strategy for accomplishing LF elimination in Haiti, and similarly resource-constrained settings, that face both programmatic and social difficulties in delivering long-term tablet-based LF MDAs.

## Methods

### Overview

We carried out a cost-revenue analysis to evaluate the production capability and financial feasibility of the developed DEC salt social enterprise via assessment of the relationship between fixed and variable costs versus the revenue received [24,27] and a modeling study centered on applying a dynamic transmission model to evaluate the cost-effectiveness of using this intervention versus standard annual MDA for eliminating LF in Léogâne arrondissement, where the salt enterprise operates [28–31]. The cost-revenue analysis was based on costs and revenue data contained in financial accounts during the production phase of the salt enterprise from 2013 to 2018, while the break-even analysis was carried out over a time horizon that ranged between 2013 to the year when the break-even point was attained. The predicted timelines to LF elimination along with the costs of annual MDA versus the net cost of supplying DEC-fortified salt until elimination was achieved were used to carry out the cost-effectiveness modeling study.

### Cost-revenue and break-even analyses

#### Cost estimation

From the operational perspective, these included periodic financial investments primarily used for purchasing large equipment; fixed costs (administrative, marketing, maintenance, and leasing costs) and variable costs (costs of raw salt, salt packaging, custom fees, labor, transportation, and drug supplies). As the salt enterprise used a product-mix business strategy of producing three different types of salt to meet demands and application in different market segments, viz. industrial - untreated salt processed to meet the requirements of various industrial applications, coarse and fine single-fortified salt (treated with 40 ppm of potassium iodate only) and double-fortified salt (treated with both 40 ppm of potassium iodate and 0.32% DEC [w/w]), to assure price competitiveness of the double-fortified salt with the local table salt in the market and to meet customer preference for either coarse or fine edible salt, we additionally estimated the variable costs of the two fortified salt types. At the project level, we multiplied the unit cost with the unit quantity of each cost item consumed on a yearly basis to obtain the annual total cost of producing all three types of salt.

#### Revenue estimation

The unit sale price was multiplied by the unit quantity sold per year to quantify the annual revenue obtained by the sale of each salt type. Total revenue was simply the sum of revenues obtained from the sale of each salt type.

### Forecasting break-even time points

This was conducted by projecting forward the total costs (investment + fixed + variable costs) and total revenues from the sale of all three types of salt estimated for different periods between 2013 to 2018 until the point at which cash flow from the project (total revenue - total cost) becomes zero or the break-even for the enterprise is attained [24–27]. Simple linear forward projections were used to make these calculations. Costs and revenues (both in US$) were used undiscounted in this study.

### Cost-effectiveness modeling

#### Overview of the LF transmission model

We extended the data-driven Monte Carlo population-based EPIFIL model for predicting local LF infections [32–36] to include comparative costs and simulations of the effectiveness of the standard two-drug (DEC plus Albendazole (ALB)) tablet-based MDA used in Haiti versus consumption of double-fortified salt sold by the salt enterprise to perform this analysis. We used a data-model assimilation technique based on the Bayesian Melding (BM) algorithm to calibrate the LF model to the microfilariae (mf) prevalence data observed in our Léogâne endemic setting [28–31], and used the localized model to simulate the impact of either intervention on timelines required to the decrease the community-level mf prevalence below the WHO-mandated elimination threshold of 1% mf [1,2]. The economic cost for carrying out annual MDA was fixed conservatively at US$0.64 per person inclusive of drug cost, given the finding that this represented the average cost of treating an individual in Haiti once initial costs stabilized [29,37], while the cost of delivering DEC-medicated salt per person per year was estimated from the difference between the costs of production and the revenues gained from sales of the salt. Simulations of cost-effectiveness were carried out by fixing the Léogâne arrondissement population at 500,000 [38]. Modeling of the impacts of MDA using DEC+ALB and DEC-medicated salt followed our previous methods published in detail in Smith et al [18]. Outcomes were compared at coverages of 65% and 80%, while effectiveness of DEC-medicated salt was also investigated at population coverages that could be obtained as salt production increased over time.

#### Input epidemiological data

The data sources used for calibrating the LF model were collected from Léogâne commune, Haiti [28–31]. The epidemiological data inputs encompassed information on baseline community-level mf prevalence (15.5%), the annual biting rate (ABR) estimated inversely by model fitting to mf prevalence data [39], and details regarding the dominant mosquito genus (*Culex quinquefasciatus*). Published details of the MDA interventions, including the relevant drug regimen, carried out in this setting during 2000-2008 were also assembled and used as required [20,28–31]. All model parameters, functions, and fitting procedures specific to this work are given in detail in S1 Supporting Information.

## Results

### Costs and production of salt

Table 1 summarizes the investment, fixed and variable costs incurred in establishing and operating the Léogâne DEC salt enterprise. These costs are presented for the years between 2013-2018 when production of food-grade fortified salt began (following an experimental phase which addressed technical issues in the fortifying of salt with DEC) along with corresponding data on the quantity of the three different types of salt (industrial, coarse and fine single-fortified, coarse and fine double-fortified) produced and sold. Note that investments occurred periodically during different expansion phases (2013, 2014, and 2017), and were primarily used to acquire capital items either from the US or from within Haiti. These were recorded as fixed assets, and comprised factory items, such as different types of pumps, screens, control systems, hoppers, sealers, storage tanks, generators, and office equipment. Fixed costs, i.e., costs that remain the same whatever the level of output produced or products sold included operating expenses, while variable costs comprised costs of items which scaled with production volume [24,25]. Examples of components of the latter two cost types incurred are given in Methods.

**Table 1.** Cost (investment, fixed cost and variable cost (in US$)) and summary of all types of salt (industrial salt, coarse and fine single-fortified salt, and coarse and fine double-fortified salt) produced (tons) from 2013-2018.

| Year | Cost (US\$) | | | Salt Production (Tons) | | | | |
| --- | --- | --- | --- | --- | --- | --- | --- | --- |
|  | Investment | Fixed Cost | Variable Cost | Industrial | Single-fortified |  | Double-fortified |  |
|  |  |  |  |  | Coarse | Fine | Coarse | Fine |
| 2013 (Jan-Dec) | 98,418 | 36,612 | 75,176 | 10 | 0 | 0 | 230 | 0 |
| 2014 (Jan-Dec) | 397,249 | 180,589 | 120,296 | 272 | 1 | 5 | 470 | 0 |
| 2015 (Jan-Dec) | 0 | 275,495 | 225,367 | 557 | 146 | 33 | 334 | 7 |
| 2016 (Jan-Dec) | 0 | 275,495 | 392,882 | 2140 | 452 | 206 | 479 | 61 |
| 2017 (Jan-Dec) | 49,280 | 558,904 | 601,308 | 2811 | 648 | 308 | 979 | 91 |
| 2018 (Jan-Dec) | 23,000 | 352,000 | 800,000 | 2431 | 1013 | 527 | 1841 | 33 |

The data on salt production show that initial output was low (e.g., 10 metric tons of industrial salt and 230 tons of coarse double-fortified salt in 2013, compared to 272, 6, and 470 tons of industrial, single-fortified, and double-fortified salt respectively in 2014). To increase production, a three phase expansion of the project was introduced beginning in the year 2013. Phase 1 focused on installing higher capacity processing equipment (January 2013-March 2014), while Phase 2 aimed to expand processing capacity by adding bulk storage and extension of the brine-washing system (May 2014-March 2015), and Phase 3 further expanded these washing, processing and storage capacities (January 2017-June 2018). These expansion phases meant that both the fixed and variable costs of salt production varied over time (Table 1), but in general, and as expected, as production expanded the variable costs increased proportionately while the fixed costs simply reflected increases in capital investments. By contrast, the production figures show that the manufacture and sale of each type of salt increased steadily, with industrial salt dominating production particularly toward the later years followed by double-fortified (i.e., DEC and iodine medicated) and single-fortified (iodine-medicated only) coarse salts. Given the population preference for coarse edible salt in Léogâne (and Haiti in general), fine salt production and sale lagged behind in volume (Table 1). The figures, however, demonstrate that with expansion in capacity the enterprise was able to achieve significant production volumes for the double-fortified salt by year 2018 (1841 metric tons recorded for the coarse variety).

### Net cash flow and break-even forecasting

The total annual costs of salt production (= Investment + Fixed Costs + Variable Costs) and revenues attained from the sale of all three types of salt are given in Table 2. Total annual costs increased as production expanded (Table 1) from US$210,206 in 2013 to US$1,175,000 in 2018 - i.e., approximately five times - but the figures show that revenue increased even faster, up to 17 times that obtained initially in 2013 (US$61,636) to close to cost of production by 2018 (US$1,064,000). The net cash flow [24–27] figures in the Table, which represent the difference between total production cost and total revenue, although being negative for all the years from 2013-2018, capture this increasing revenue returns (as reflected by the declining negative trend in the net cash flow) towards the later years, suggesting that the project is close to achieving break-even or profitability in the near future. To estimate the exact time point when the enterprise is likely to break-even (i.e., the time point when the total program cost is equal to the total revenue), we employed a simple linear model to project forward the total project costs and revenues calculated for 2013-2018. Fig 1(a) shows that if we use the full data on annual costs and revenues obtained for the whole 2013-2018 period, the project will break-even in 2027. However, if we use the data from 2016-2018, when total salt production had reached significant levels (Table 1), to predict the time point at which the break-even point will be achieved, this will occur earlier by year 2022 (Fig 1(b)).

**Fig 1.**
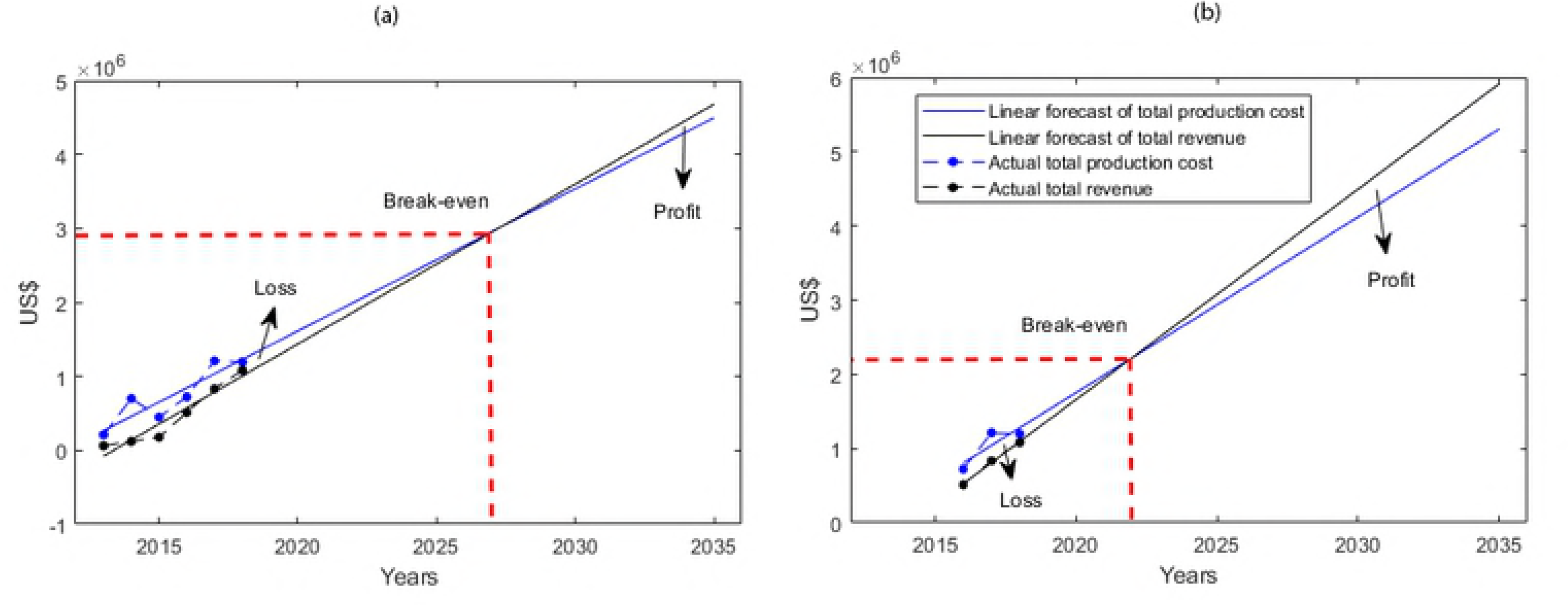
Forecasts of the break-even years of the salt enterprise. Results are shown from using a simple linear model to project forward the total project costs and revenues (in US$). **(a)** Forecast of break-even year using total cost and total revenue data calculated for 2013-2018 given in Table 2. **(b)** Forecast of break-even year using total cost and total revenue data calculated for 2016-2018 given in Table 2. The red dotted line indicates the break-even point for each scenario.

**Table 2.** Year wise details of total cost, total revenue and net cash flow attained by the Haiti salt enterprise between 2013 to 2018. The total costs shown are calculated using the cost data for the production of all three types of salt, while total revenue is calculated using the revenue data from the sale of all three types of salt for each year.

| Year | Total Cost (US\$) | Total Revenue (US\$) | Net Cash Flow |
| --- | --- | --- | --- |
| 2013 | 210,206 | 61,636 | -148,570 |
| 2014 | 698,134 | 118,583 | -579,551 |
| 2015 | 448,168 | 175,629 | -272,539 |
| 2016 | 721,070 | 512,588 | -208,482 |
| 2017 | 1,209,492 | 832,285 | -377,207 |
| 2018 | 1,175,000 | 1,064,000 | -111,000 |

Fig 2 depicts the per ton revenues from sales and costs of producing the three different types of salts for the years 2013-2018. The results show that for all salt types, while initially there was a large difference between the production cost and revenue per ton (i.e., a large negative cash-flow), this difference decreased for each salt category with time. This occurred faster, however, in the case of the industrial and single-fortified salt, such that break-even was achieved by 2018. By contrast, the cash-flow from the production and sale of the double-fortified salt was still negative (lower revenue compared to production costs) at the end of the present study period of 2018. This result shows, first, that the overall break-even estimated in this study (Fig 1) for the project is due to the delay in reaching the break-even year for double-fortified salt. Second, it also highlights how the mixed product strategy of producing different types of salt targeting different market sectors can allow the more profitable products (industrial, single-fortified salt) to subsidize the sale of a product (double-fortified salt) whose cost (approximately US$200 per ton (Fig 2)) needs to be kept competitive with other edible salt sold (retailed at $US265 per ton in Haiti (James Reimer, personal communication)) in the local market.

**Fig 2.**
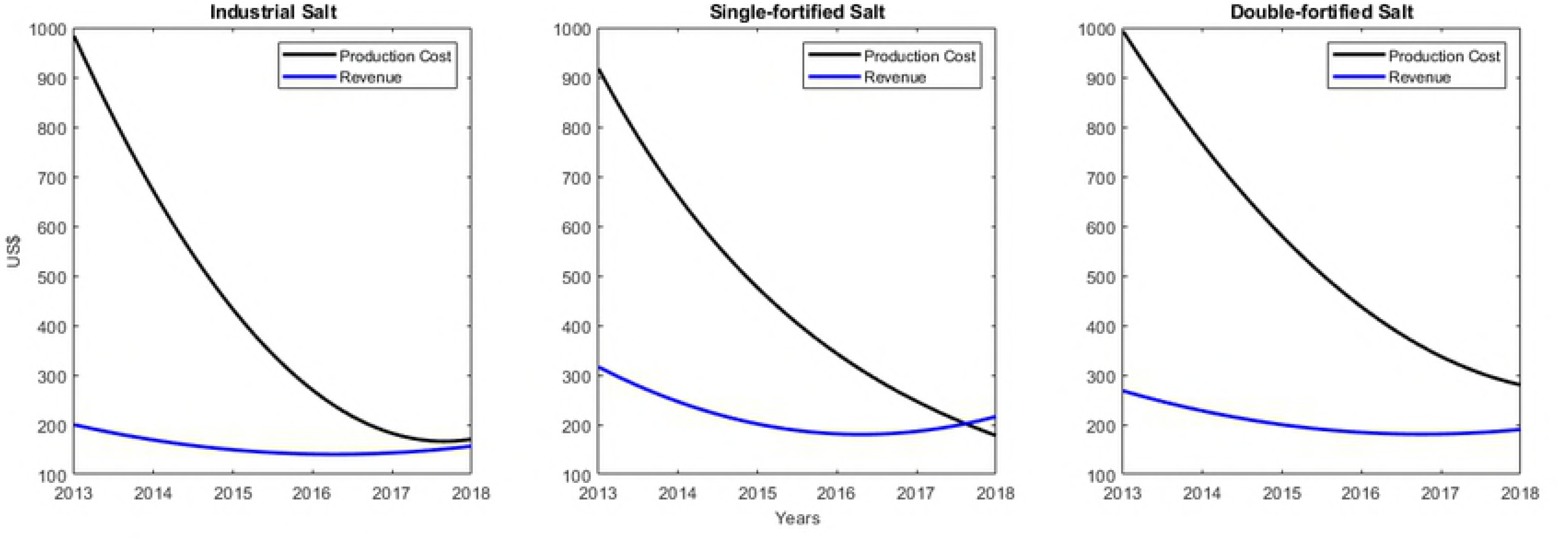
Revenue and production cost for industrial, single-fortified and double-fortified salt. Both revenue and production cost (in US$) for each salt are shown per ton.

### Marginal cost of manufacturing DEC-medicated salt

This was evaluated by analysis of the difference in the variable costs of producing the single-fortified (which included only potassium iodate) versus the double-fortified (both potassium iodate and DEC included) salt types, given that the investment and fixed costs going into manufacturing all salt types were shared equally between each type. The variable costs per ton for producing the single-fortified (coarse and fine) and double-fortified (coarse and fine) salt for two types of bags/bales (25.0-kg bags and 12.5-kg bales) are provided in Table 3 for the years 2014-2018 when both salt products were produced (note that production of single-fortified salt began only in 2014 (Table 1)). Analysis of the difference in the variable costs for producing the single-fortified versus the double-fortified salt indicate that this was consistently about US$70 per metric ton, irrespective of which type - coarse or fine variety - or types of bags/bales were produced (Table 3). Given that the cost of DEC, as delivered by Syntholab Chemicals to the project was US$21.60 per kg (James Reimer, personal communication), and DEC salt in this project was fortified with 0.32% DEC by weight or with 3.2kg of DEC/ton, it can be seen that the cost of producing one metric ton of DEC salt works to be US$ 69.12/ton (i.e., 3.2 kg DEC x US$21.60 per kg). This result indicates that the marginal cost, or difference in the variable cost, of adding DEC to single-fortified salt (US$70; Table 3) was simply due to the purchase cost of DEC.

**Table 3.** Marginal cost of manufacturing DEC-medicated salt. This was evaluated by analysis of the difference in the variable costs of producing the single-fortified (which included only potassium iodate) versus the double-fortified (both potassium iodate and DEC included) salt types, given that the investment and fixed costs going into manufacturing all salt types were shared equally between each type.

| Year | Types of Bags/Bales | Variable cost of Coarse Salt (US\$/Ton) | | | Variable cost of Fine Salt (US\$/Ton) | | |
| --- | --- | --- | --- | --- | --- | --- | --- |
|  |  | Single-fortified | Double-fortified | Difference | Single-fortified | Double-fortified | Difference |
| 2014 | 25.0-kg Bags | 150 | 220 | 70 | 160 | 230 | 70 |
|  | 12.5-kg Bales | 190 | 260 | 70 | 200 | 270 | 70 |
| 2015 | 25.0-kg Bags | 145 | 215 | 70 | 155 | 225 | 70 |
|  | 12.5-kg Bales | 185 | 255 | 70 | 195 | 265 | 70 |
| 2016 | 25.0-kg Bags | 145 | 215 | 70 | 155 | 225 | 70 |
|  | 12.5-kg Bales | 185 | 255 | 70 | 195 | 165 | 70 |
| 2017 | 25.0-kg Bags | 135 | 205 | 70 | 145 | 215 | 70 |
|  | 12.5-kg Bales | 175 | 245 | 70 | 185 | 255 | 70 |
| 2018 | 25.0-kg Bags | 135 | 205 | 70 | 145 | 215 | 70 |
|  | 12.5-kg Bales | 175 | 245 | 70 | 185 | 255 | 70 |

### Population coverage attained by the DEC salt enterprise

Table 4 shows the potential increase in demand for salt in the Léogâne arrondissement calculated as a function of changes in population size from 2013 to 2018. It also presents the potential DEC-fortified salt coverage which may be achieved in the setting by increasing sale of the double-fortified salt produced over this period. The annual population size estimates from 2013 onwards were predicted using a growth rate of 1.28% [38], whereas the yearly population demand for edible salt was calculated by assuming that the daily salt consumption per person is 15gm (average of the reported daily per-capita consumption in Haiti of 10gm and 19gm [5,40]). Assuming that the sale of double-fortified salt is widespread in the community (i.e., not targeted towards one segment of the population), coverage of DEC salt in Léogâne can then be roughly estimated simply by dividing the quantity of salt produced over the estimated demand. The results from this calculation, listed in the last column of Table 4, shows that as production increased rapidly from 2013 to 2018, this could increase potential population coverage achieved by sales of the DEC-medicated salt from as low as 8.45% in 2013 to approximately 65% in 2018.

**Table 4.** Details of production of double-fortified (DEC-medicated) salt (tons), its potential demand/year, and coverage in Léogâne Arrondissement.

| Year | Quantity of double-fortified (DEC-medicated) salt (Tons) | Population of Léogâne Arrondissement* | Potential Demand of salt/year (Tons)** | Coverage (%) |
| --- | --- | --- | --- | --- |
| 2013 | 230 | 497,274 | 2,723 | 8.45 |
| 2014 | 470 | 503,241 | 2,755 | 17.06 |
| 2015 | 341 | 509,280 | 2,788 | 12.23 |
| 2016 | 540 | 515,799 | 2,824 | 19.12 |
| 2017 | 1070 | 522,401 | 2,860 | 37.41 |
| 2018 | 1874 | 529,088 | 2,897 | 64.69 |

\*Population growth rate of Haiti is 1.28% [38]
\*\*Assuming that the salt consumption rate per person per day is 15gm [5,40]

### Cost-effectiveness modeling

This was carried out by comparing the costs and effectiveness achieved by the production and sale of DEC-fortified salt versus implementation of the annual tablet-based DEC/ALB MDA for accomplishing LF elimination in Léogâne (as measured by the times by which the initially observed baseline pre-control mf prevalence in this setting reach below the WHO-recommended 1% mf prevalence threshold [1,2]). Timelines taken by each intervention to reduce the initial prevalence to below the 1% mf threshold were quantified by simulating trends in infection prevalence due to these interventions using the Monte Carlo-based EPIFIL model localized to reflect LF transmission in Léogâne [32–36]. The impact of the annual tablet-based MDAs was studied by running the model at 65% and 80% population coverages, while we simulated the effect of the produced DEC salt using 65%, 80% and the actual population coverages given in Table 4. EPIFIL was calibrated to the pre-control mf prevalence data observed in Léogâne (15.5% [28]) so that it could reproduce the baseline age-infection prevalence for carrying out this analysis.

Fig 3 portrays the predicted timelines to LF elimination in Léogâne under each of the MDA and salt interventions investigated. For MDA, the depicted simulations indicate that it would take up to 7 years at 65% coverage, and 5 years at 80% coverage, respectively, to reach the 1% mf threshold. By contrast, the model predictions show that it will take just 1 year at 65% coverage, 5 months at 80% coverage, and 3 years if actual population coverage (Table 4) are used to reduce the pre-control prevalence to below this threshold (Fig 3). This highlights the dramatic effect that daily consumption of DEC-medicated salt even at low dosages (0.32% w/w) would have compared to annual intake of a higher dosages of DEC (and ALB) as provided by tablet-based MDA for eliminating LF infection in an endemic setting [12,15–18]. Note that the actual DEC salt population coverages (Table 4) used in this analysis assume that salt supply occurred uniformly and sale was restricted to Léogâne arrondissement only. Any changes in these parameters would mean attaining lower annual population coverages than shown in Table 4; however, a sensitivity analysis using coverage values 15-20% lower than those depicted in the Table did not affect the above timelines significantly (data not shown).

**Fig 3.**
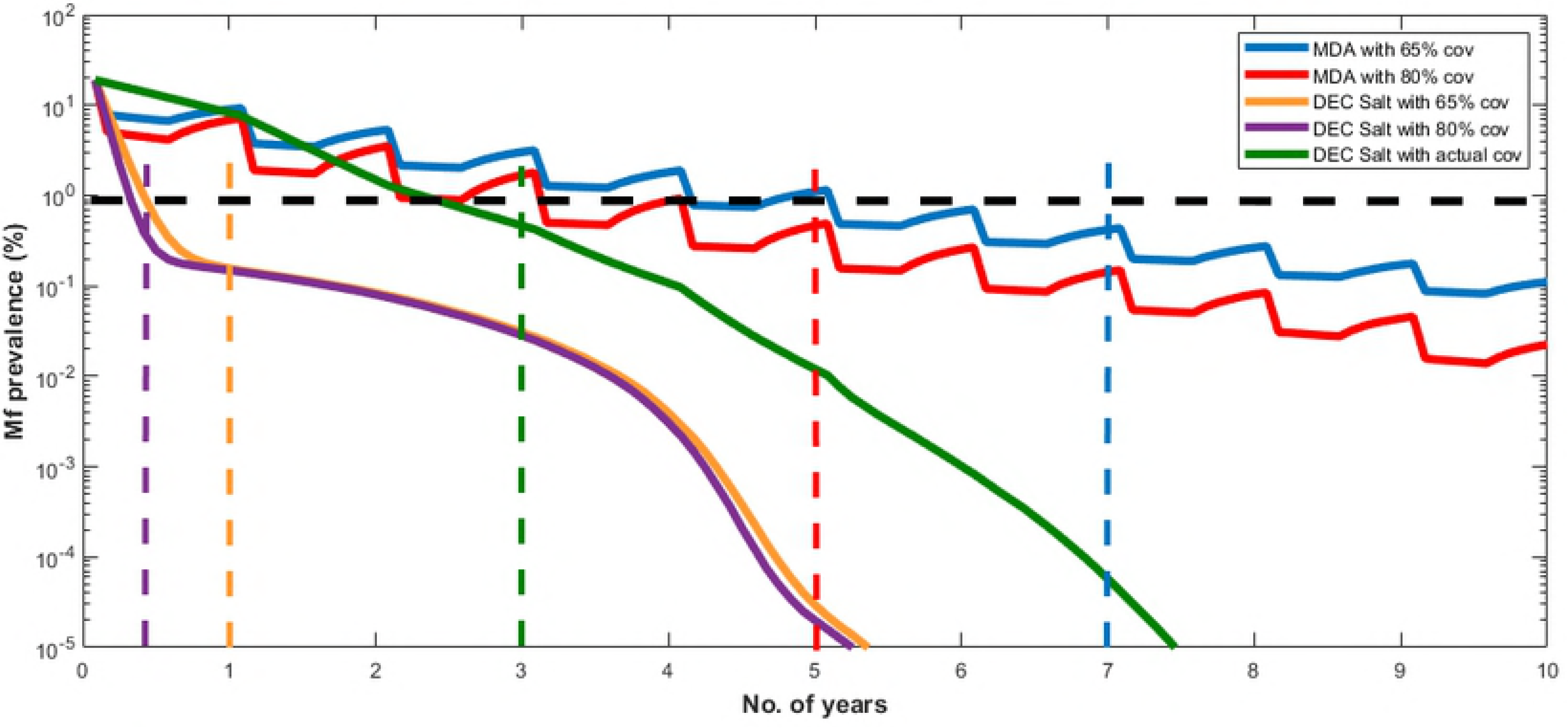
Timelines to reach the WHO recommended 1% mf threshold for MDA implanted at 65% and 80% coverage and DEC-medicated salt at 65%, 80%, and actual population coverages. The median model predicted trajectory is shown for each intervention strategy. The black dotted horizontal line indicates WHO recommended 1% mf prevalence threshold. The blue, red, orange, purple and green dotted vertical lines indicate the time points to reach 1% mf threshold using either MDA at 65% and 80% coverages or DEC Salt at 65%, 80%, and actual coverage respectively.

The comparative costs of carrying out annual MDA versus supplying DEC-medicated salt are shown in Table 5. These show that while the total cost of delivering annual MDA (here fixed at US$0.64 per person, inclusive of the cost of the drug [29,37]), simply scaled with population growth, and will continue to be substantial on a yearly basis until LF elimination is achieved, the net cost of supplying DEC-fortified salt through the social business model will decline dramatically with time as production costs decrease and revenues begin to increase over time (Fig 2). Indeed, projection forward of the net or revenue - production cost data collected during the years 2013-2018 indicates that the total and per capita net costs of DEC salt provided through the present social enterprise could potentially even become zero at the time point (2027) when the project breaks even (Fig 1(a)).

**Table 5.**
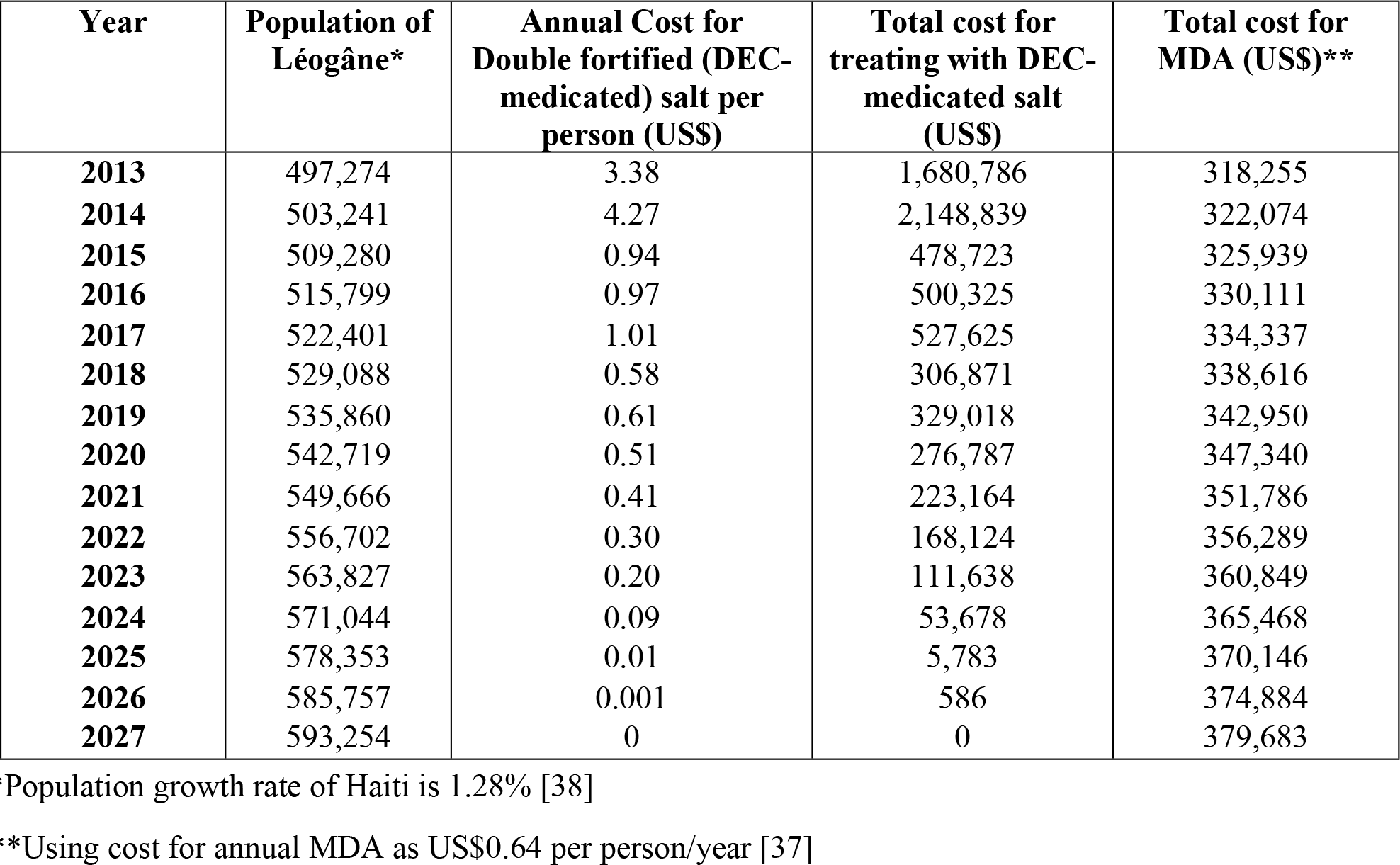
Forecast of the population of Léogâne, annual cost of supplying double-fortified (DEC-medicated) salt per person per year, and total costs of treatment with DEC-medicated salt and MDA respectively for the years 2013-2027. The population growth rate for Haiti estimated in year 2015 (1.28%) was used in making the population size estimations shown in the Table, whereas a simple linear model based on the cost and revenue data connected with DEC-medicated salt production and sale (both fine- and coarse salt types) for the years 2013-2018 (Fig 2) was used to forecast the annual cost of salt-based DEC treatments per person and for the total population between 2019-2027.

| Year | Population of Léogâne* | Annual Cost for Double fortified (DEC-medicated) salt per person (US\$) | Total cost for treating with DEC-medicated salt (US\$) | Total cost for MDA (US\$)** |
| --- | --- | --- | --- | --- |
| 2013 | 497,274 | 3.38 | 1,680,786 | 318,255 |
| 2014 | 503,241 | 4.27 | 2,148,839 | 322,074 |
| 2015 | 509,280 | 0.94 | 478,723 | 325,939 |
| 2016 | 515,799 | 0.97 | 500,325 | 330,111 |
| 2017 | 522,401 | 1.01 | 527,625 | 334,337 |
| 2018 | 529,088 | 0.58 | 306,871 | 338,616 |
| 2019 | 535,860 | 0.61 | 329,018 | 342,950 |
| 2020 | 542,719 | 0.51 | 276,787 | 347,340 |
| 2021 | 549,666 | 0.41 | 223,164 | 351,786 |
| 2022 | 556,702 | 0.30 | 168,124 | 356,289 |
| 2023 | 563,827 | 0.20 | 111,638 | 360,849 |
| 2024 | 571,044 | 0.09 | 53,678 | 365,468 |
| 2025 | 578,353 | 0.01 | 5,783 | 370,146 |
| 2026 | 585,757 | 0.001 | 586 | 374,884 |
| 2027 | 593,254 | 0 | 0 | 379,683 |

\*Population growth rate of Haiti is 1.28% [38]
\*\*Using cost for annual MDA as US\$0.64 per person/year [37]

We used the total costs of implementing each strategy until LF elimination (crossing below the 1% mf threshold) is achieved to investigate the cost effectiveness of either strategy. The total costs and effectiveness of using MDA at 65% and 80% coverages and supplying DEC salt at 65%, 80% and the actual coverages given in Table 4, were evaluated and compared via calculations of the average cost effectiveness ratio [10,41–47]. With respect to DEC salt, we also conducted the analysis using three different net costs of salt production per person per year as recorded in Table 5: (i) average of the net cost of salt produced per person calculated for the years 2018-2020, i.e., US$0.57 per person/year, (ii) average of the net cost of salt per person for the years 2021-2023, i.e., US$0.3 per person/year, and (iii) average of the net cost of salt per person for the years 2025-2027, i.e., US$0.025 per person/year. This was performed to assess the sensitivity of the present results to changes in the steeply declining net cost of salt production over time observed in this study.

The results from this exercise are shown in Table 6, and indicate, principally, that irrespective of coverage, the costs of using MDA are significantly greater than those arising from using the salt strategies, primarily because of its lesser effectiveness as well as higher and stationary unit cost. Among the three salt scenarios, costs for eliminating LF, as expected, declined with decreasing net cost of production over time with scenario three showing the lowest costs. However, for all strategies, while the most effective strategy is to deliver MDA or salt at 80% coverage, the most cost-effective option (in terms of minimizing the cost-effectiveness ratio) occurred at either 65% drug coverage for MDA or at the lower actual coverages recorded for DEC salt (Table 6). This suggests the existence of a trade-off between cost and effectiveness (in terms of duration of interventions required to achieve elimination) in the case of these strategies, i.e., obtaining a higher coverage for accomplishing earlier LF elimination comes with heightened costs. Overall, however, the results indicate that because of: 1) decreasing net cost of DEC salt production over time, and 2) expansion of its coverage in the population leading to significantly reduced elimination timelines, the delivery of DEC through a salt enterprise may be significantly more cost-effective than annual DEC tablet-based MDA for accomplishing LF elimination in the Léogâne arrondissement setting. It is also to be appreciated that we used a conservative treatment cost of $0.64 per person for modeling the cost-effectiveness of the tablet-based MDA program in this study. This represented a best-case scenario for the MDA program implemented in Léogâne given that the actual economic costs, i.e., inclusive of the cost of donated drugs, started out higher (US$1.84 over the first 3 MDAs) before approaching the stabilized value used in our analysis as the program became more efficient [29,37]. Indeed, a recent systematic review indicated that MDA program costs can vary substantially between settings, with an average economic cost that could reach as high as US$1.32 [48]. Use of such values or inclusion of the actual change observed in the per person treatment cost over time in Léogâne in the present analysis would clearly further increase the cost of MDA over that presented in Table 5, which in turn would lead to an even higher cost-effectiveness ratio for MDA compared to those estimated for DEC salt in this study (Table 6).

**Table 6.** Comparison of total costs and CE ratios of population treatment using MDA versus supply of DEC-medicated salt for different drug coverages and DEC salt costs.

| **Treatment Method** | **Coverage** | **No. of years to reach 1% mf threshold** | **Total Cost (US\$)** | **CE ratio** |
| --- | --- | --- | --- | --- |
| **Annual MDA\*** | 65% | 7.00 | 1,456,000 | 208,000 |
|  | 80% | 5.00 | 1,280,000 | 256,000 |
| **DEC salt<sup>1</sup>** | 65% | 1.00 | 185,250 | 185,250 |
|  | 80% | 0.42 | 95,760 | 228,000 |
|  | Actual<sup>†</sup> | 3.00 | 105,450 | 35,150 |
| **DEC salt<sup>2</sup>** | 65% | 1.00 | 97,500 | 97,500 |
|  | 80% | 0.42 | 50,400 | 120,000 |
|  | Actual<sup>†</sup> | 3.00 | 55,500 | 18,500 |
| **DEC salt<sup>3</sup>** | 65% | 1.00 | 8,125 | 8,125 |
|  | 80% | 0.42 | 4,200 | 10,000 |
|  | Actual<sup>†</sup> | 3.00 | 4,625 | 1,542 |

\*Using cost for annual MDA as US\$0.64 per person/year [37], and total population of Léogâne arrondissement 500,000 (fixed).
<sup>†</sup>Actual coverage of DEC-medicated salt given in Table 4.
<sup>1</sup>Average of the net cost of DEC salt produced per person calculated for the years 2018-2020 given in Table 5, i.e., US\$0.57 per person/year.
<sup>2</sup>Average of the net cost of DEC salt produced per person calculated for the years 2021-2023 given in Table 5, i.e., US\$0.3 per person/year.
<sup>3</sup>Average of the net cost of DEC salt produced per person calculated for the years 2025-2027 given in Table 5, i.e., US\$0.025 per person/year.

## Discussion

In this study, we have undertaken a performance assessment of a novel social enterprise developed through a collaboration between Haitian and international partners engaged with LF control in the country, as a means to enhance the delivery of anti-filarial drugs to populations through the trading of salt co-fortified with DEC. Although DEC-fortified salt has been used previously in both pilot and region-wide LF intervention programs in a variety of global regions, ranging from Brazil, Tanzania, India and China to effectively control or eliminate LF [5,12–17], it is to be noted that the developed Haiti salt enterprise is the first attempt anywhere in the world to apply the principles of social entrepreneurship for delivering such an intervention. Recent work has highlighted how such social enterprises - that is, a social mission-driven organization that trades in goods or services for a social purpose - are emerging as a potentially effective supply side solution to the provision of cost-efficient public services in response to government failures, business that seek to extract maximal returns on investment, and unstable non-profit organizations [21–23]. In particular, this work has shown how these business entities can solve social problems via their potential to deliver greater responsiveness, efficiency and cost-effectiveness, through an explicit focus on meeting specific social goals while operating with the financial discipline and innovation of a private-sector business [21–23,49,50].

Although there is continuing debate as to how best to evaluate the performance of social enterprises, it is clear that at least two basic components related to the bottom line of these entities require assessment [21,23]. These primarily include: the economic-financial component for measuring overall organizational efficiency, profitability and hence sustainability, and the social effectiveness of the enterprise [21–23]. Here, we have combined the tools of financial accounting and modeling of cost-effectiveness to measure these components in order to present a first analysis of the utility of using the developed Haitian DEC salt enterprise as a sustainable and economically efficient strategy to bring about LF elimination in programmatically difficult-to-control settings, like the arrondissement of Léogâne, Haiti [10,41–47].

Our analysis of the performance of the present salt enterprise for creating social value first focused on the question of capacity to economically produce sufficient amounts of DEC-fortified salt for significantly affecting the elimination of LF in the study setting. The production figures shown in Table 1 indicate, firstly, that while initial production of all types of salt were low during the initial years of operation, by 2018, and just 5 years after processing began in 2013, the enterprise had reached high levels of both total (5,845 metric tons) and DEC-fortified salt (1,841 metric tons) production, respectively (Table 1). Our analysis of the population coverage that the sale of DEC salt could provide demonstrate that the amounts produced could have potentially resulted in a coverage rate of 65% by 2018 (Table 4), which our previous modeling study [18] and the present cost-effectiveness exercise (Fig 3 and Table 6) indicate is sufficient to accelerate the achievement of LF elimination in the Léogâne setting. These findings suggest that as the result of the expansion phases carried out through new capital investments (Table 1), the current DEC salt project may have reached the production capacity required to achieve its stated social mission of using a market-based approach for delivering sufficient edible DEC-medicated salt as a means to bring about efficient LF transmission interruption in the present study setting.

Assessment of the economic and financial performance of the salt enterprise carried out in this study using cost-revenue analysis and financial forecasting has provided further insights regarding the organization efforts used to reach economic equilibrium and hence trading viability. This is an important consideration for evaluating the performance of social enterprises because first and foremost these entities are enterprises, and therefore their social goals can be pursued only by ensuring economic and financial fidelity [21–23]. Our major result in this area is providing clarity regarding how the enterprise’s achieved outputs in salt production and sale may affect its potential to reach break-even points (Fig 1). Specifically, we show that given the observed trends in production costs and revenue to 2018 (Table 2), the present salt enterprise may either break-even by 2027 (if we forecast linearly using all the data from 2013 to 2018), or as early as by 2022 (if we use data collected during 2016-2018 after the project had significantly expanded capacity). This result is clearly dependent on assuming that capacity to produce the increased amount of salt to meet either break-even points is available within the enterprise without any further expansion, and demand for the produced salt in all sectors (industrial to edible salt markets) will also expand commensurately. Nonetheless, the finding that it might be possible to reach the break-even point by 2022 (i.e., over the next 4 years) is encouraging, and suggests that the enterprise is likely to be self-sustaining and could become profitable in the very near future. Indeed, analysis of trends in costs of production and revenues gained per ton of each category of salt (Fig 2) indicate that both the industrial and single-fortified salt categories may already have reached their individual break-even points in 2018, and that the delay for achieving break-even status by the social enterprise is primarily due to the lag experienced by the production and sale of the double-fortified salt. Although the per ton production cost of the latter salt declined as significantly over time as the other two salt categories (Fig 2), indicating the achievement of considerable economics of scale, the need to keep the price of the DEC salt below the marginal cost of adding the drug ($70/metric ton (see Table 3)) to compete with untreated local edible salt in the market means that either: 1) the current market price needs to be revised upwards, or 2) further economies of scale need to be found to bring down production costs, or 3) cross-subsidy from the more profitable categories of salt produced will be required in order to continue with the processing of DEC salt in this setting. While on the one hand, such a capacity to use a product mix strategy innovatively as a means to subsidize the marketing of a product for meeting a social need is a feature of using an enterprise model, note that this may be a particular effect of developing markets in settings, such as Léogâne and Haiti in general, where a strong market-based economy is only just evolving. For other LF endemic settings with stronger market economies and established salt industries, the need for such subsides may be significantly lower meaning that the sale of DEC salt could occur at nearer the true marginal cost of production, i.e., at the actual cost of purchasing the drug itself. Note also that our present forecasts do not fully consider the likely impacts of key swing factors that may significantly affect the profitability of the salt social enterprise, such as enforcement of the 2017 law requiring all food salt in Haiti to be fortified, further progress on market segmentation and the resulting product mix, and significant weather events similar to Hurricane Michael in 2016. Positive changes in the first two factors will clearly enhance the enterprise’s ability to break-even faster and hence attain profitability sooner than predicted in this work.

The cost-effectiveness modeling exercise carried out in this study showed that apart from the efficiency of the business model used for achieving economic and financial sustainability, the salt enterprise may also be more cost-effective than the standard tablet-based annual MDA program for accomplishing LF elimination in Léogâne (Table 6). This is because not only will the population coverage that can be potentially attained by sale of the DEC salt (Table 4) be sufficient to make this strategy more effective than annual MDA in reducing the number of years (3 versus 7 years) required for achieving LF elimination (as defined by reducing mf prevalence below the WHO threshold of 1% [2]), the total overall costs involved - due to both decreasing net cost of production and the need for shorter durations of control - for using the salt approach are also significantly lower than those which will be incurred in running the MDA program. Indeed, this greater social impact of using the present social salt enterprise compared to annual MDA was found to be a general outcome, irrespective of the other intervention coverages investigated (i.e., at 65% coverage - the often normal coverage obtained by MDA programs - or at the recommended optimal coverage of 80% [2]) (Table 6). These results add to our recent modeling work, which highlighted how the continuous consumption of the drug, even at low daily per capita dosages, by resulting in a cumulative impact on the survival of worms and mf which is significantly higher than that afforded by the higher-dosed annual MDA treatment, make DEC medicated interventions, even when delivered at moderate population coverages, a markedly potent strategy for interrupting LF transmission [18]. Finally, an intriguing possibility highlighted by the break-even analysis and the cost forecasting results shown in Fig 1 and Table 5 is that using a social enterprise strategy for delivering DEC through marketing of medicated salt could in principle also lead to zero disease elimination cost for the community once the social business attains profitability (i.e., return a positive cash-flow). This is an important result, and demonstrates how using a social enterprise that pursues a social goal by production of services and goods whilst respecting economic efficiency may offer an effective, financially sustainable, intervention strategy in settings facing major fiscal, infrastructural and logistical barriers to carrying out tablet-based programs aiming to control or eliminate parasitic infections.

The present performance evaluation primarily focused on internal (labor, capital, income and taxes) and external (goods and services bought outside the company) expenses/resources related to the economic viability of the salt enterprise [21]. However, estimation of the full social value of a sustainable health social enterprise must also consider, apart from the social benefits accruing from reducing disease only, the wider consequences for a community [21]. Benefits here could be via the choice and use of resources that further address the community interest, such as choosing local salt suppliers to favor short supply chains, choosing socially certified suppliers, adopting a regime of decent work conditions and even giving employment to workers coming from disadvantaged backgrounds [21]. Such analysis must also include calculation of the larger social benefit associated with the potential for the double-fortified salt to additionally and simultaneously reduce the impacts of iodine deficiency in the population [5]. Note, additionally, that the present salt enterprise represents the first attempt to build industrial-scale capacity on the island for processing large volumes of salt to meet various local needs, which apart from providing a market for local raw salt producers can also act as means to significantly stabilize the price of salt sold in the local markets. These benefits, however, must be contrasted against potential adverse effects, such as domination of the market by the growing enterprise, requiring an analysis of how best to compensate for such loses. Recent developments in applying Social Returns on Investment (SROI) approaches for comparing the full monetized social costs of a program with the full monetized social benefits of achieving a health outcome (or set of outcomes) may offer a means for undertaking this fuller analysis [22,51].

We have also used rough first calculations of the population coverages that could be obtained with the expansion of salt production in the present cost-effectiveness modelling study. Field studies to assess the actual household coverage achieved through the enterprise will be critical for not only more realistically quantifying its effectiveness for accomplishing LF transmission interruption in a community, but also for identifying better marketing strategies to achieve good population coverage.

In conclusion, we have presented an economic and financial analysis of the Haitian salt social enterprise, which indicates that it may present a sustainable and socially-responsible strategy for aiding the elimination of LF via the marketing of DEC-medicated salt in settings facing fiscal, infrastructural and logistical challenges for delivering tablet-based elimination programs. Results from the break-even projections carried out in this study indicate that the strategy may even have the potential to achieve zero societal costs once it attains profitability (i.e., results in a positive cash-flow). This study further has shown that the Haitian salt enterprise may have already reached production and sales levels that could result in the coverage of the Léogâne study population at proportions sufficient enough to break LF transmission. Finally, our simulation-based cost-effectiveness study has indicated that because of: 1) increasing revenue from the sale of the DEC salt obtained over time, 2) expansion of its delivery in the population, and 3) the effect of continuous consumption of the drug, even at low daily per capita dosages, leading to a cumulative impact on the survival of worms and mf higher than that afforded by the higher-dosed annual MDA treatment [18], the delivery of DEC through the present Haiti salt enterprise may represent a significantly more cost-effective option than annual DEC tablet-based MDA for accomplishing LF elimination. While these are encouraging first results and highlight both the economic viability and social effectiveness of using a salt enterprise in the fight against LF, it is clear that efforts to more fully quantify the social value and strategies for developing similar salt social enterprises elsewhere in other endemic settings with different market structures than those of Haiti are now required if the comparative or joint utility of the approach among the current arsenal of LF intervention strategies is to be fully appraised and understood. We note that the means by which the global iodization of edible salt has been accomplished successfully over the past two decades may offer a particularly apt model for building and sustaining the present intervention globally, and suggest that similar tactics used in that program based on introducing DEC medication into prevailing salt production and distribution systems, collaboration with the national and regional salt industries, and engagement with the government sector, civic society and the general public [52], could also make the universal deployment of DEC-medicated salt eminently possible. With less than three years remaining for meeting the 2020 target set by WHO for accomplishing the global elimination of LF, the present results indicate that these appraisals and development of policies and strategies for delivery of DEC-salt, either via deployment of similarly-fashioned salt enterprises, such as the present, or through mobilization of existing salt industries, perhaps along with health system-led MDA and vector-control programs, in socially-challenging environments, like Haiti, would improve our current efforts for meeting this laudable but exacting goal successfully.

## Acknowledgements

The authors gratefully acknowledge the support of this work by the Notre Dame Haiti Program. The authors are also thankful to Dr. Kirk Doran, Associate Professor in the Department of Economics, University of Notre Dame, for his valuable suggestions and help with the economic analyses carried out in this work. A part of the model runs was carried out using the MATLAB Parallel Computing Toolbox available on Computer Clusters of the University of Notre Dame’s Center for Research Computation.

